# Subthreshold repetitive transcranial magnetic stimulation drives structural synaptic plasticity in the young and aged motor cortex

**DOI:** 10.1101/2021.03.10.434706

**Authors:** Alexander D Tang, William Bennett, Aidan D Bindoff, Samuel Bolland, Jessica Collins, Ross C Langley, Michael I Garry, Jeffery J Summers, Mark R Hinder, Jennifer Rodger, Alison J Canty

## Abstract

**Background:** Repetitive transcranial magnetic stimulation (rTMS) is a non-invasive tool commonly used to drive neural plasticity in the young adult and aged brain. Recent data from mouse models have shown that even at subthreshold intensities (0.12 Tesla), rTMS can drive neuronal and glial plasticity in the motor cortex. However, the physiological mechanisms underlying subthreshold rTMS induced plasticity and whether these are altered with normal ageing are unclear.

**Objective:** To assess the effect of subthreshold rTMS, using the intermittent theta burst stimulation (iTBS) protocol on structural synaptic plasticity in the mouse motor cortex of young and aged mice.

**Methods:** Longitudinal *in vivo* 2-photon microscopy was used to measure changes to the structural plasticity of pyramidal neuron dendritic spines in the motor cortex following a single train of subthreshold rTMS (in young adult and aged animals) or the same rTMS train administered on 4 consecutive days (in young adult animals only). Data were analysed with Bayesian hierarchical generalized linear regression models and interpreted with the aid of Bayes Factors (BF).

**Results:** We found strong evidence (BF>10) that subthreshold rTMS altered the rate of dendritic spine losses and gains, dependent on the number of stimulation sessions and that a single session of subthreshold rTMS was effective in driving structural synaptic plasticity in both young adult and aged mice.

**Conclusion:** These findings provide further evidence that rTMS drives synaptic plasticity in the brain and uncovers structural synaptic plasticity as a key mechanism of subthreshold rTMS induced plasticity.

## Introduction

For over three decades, repetitive transcranial magnetic stimulation (rTMS) has been used to drive neural plasticity in the human brain. Traditionally, this has been achieved by delivering transient pulses of magnetic fields over the scalp at high stimulation intensities (>1 Tesla) to drive activity within neural networks (i.e., suprathreshold stimulation). However, using rodent models, we have shown that even at subthreshold intensities (e.g. 0.12 Tesla) that do not directly induce neuronal activity, rTMS can still induce neural plasticity. For example, we have shown that a single session of subthreshold rTMS, alters neuronal excitability at the single cell (layer 5 pyramidal neurons) [1] and network level (motor evoked potentials) [2] whereas multiple sessions of subthreshold rTMS have been shown to promote neuronal (e.g. skilled motor learning [3]) and glial plasticity [4, 5]. However, the effect of subthreshold rTMS on synaptic plasticity has not, to date, been explored.

Irrespective of intensity, synaptic plasticity has long thought to be the main physiological mechanism underlying rTMS induced plasticity given the ability of rTMS to alter learning and motor evoked potential amplitudes in humans and rodents [6, 7]. In addition, synaptic plasticity is impaired in the aged brain [8-10] which in part, may explain the reports of reduced rTMS-induced plasticity in older adults [11-13]. However, despite its widespread use, the physiological mechanisms underlying rTMS induced neural plasticity remain unclear, making it difficult to determine which neurological diseases or disorders are suited for rTMS intervention. While suprathreshold repetitive magnetic stimulation has been shown to induce synaptic plasticity *in vitro* (organotypic hippocampal slices) [14-16], it is unknown whether either intensity of rTMS induces synaptic plasticity *in vivo* and whether this is decreased in the aged brain.

Rodent rTMS models are a useful adjunct to human studies as they allow for direct measures of neural plasticity, including measurements of synaptic plasticity and connectivity in the living brain with longitudinal *in vivo* microscopy. Using this technique, structural plasticity of dendritic spines (the post-synaptic structure) can be quantified through changes to the density of dendritic spines and to the rates of dendritic spine gains and losses. For example, in the motor cortex, structural synaptic plasticity is a fundamental process that facilitates the learning of skilled motor behaviours, with increases to the rate of dendritic spine gains and losses on the apical dendrites of layer 5 pyramidal neurons during the learning period [17, 18]. Using a similar longitudinal *in vivo* imaging approach, here we characterised the changes to structural synaptic plasticity in the motor cortex following a single session of subthreshold rTMS in young adult and “aged” mice. Additionally, we investigated whether the effects of subthreshold rTMS are cumulative by comparing the effects of a single day of rTMS to that of multiple days of stimulation in young adult mice.

## Materials and methods

### Animals

Thy1-GFP-M mice (maintained on a C57BL/6J background) were used throughout, which express enhanced green fluorescent protein (EGFP) in a sparse subset of cortical excitatory neurons in layers 2/3 and 5 [19]. Mice were group housed on a 12-hour light/dark cycle and given food and water *ad libitum*. All animal experimentation was performed under the guidelines stipulated by the University of Tasmania Animal Ethics Committee (approval number: A0013168), which is in accordance with the Australian code of practice for the care and use of animals for scientific purposes.

Mice were classified as “young adult” aged 3-6 months, or “aged” at 23-24 months. The single session of rTMS group (total n=10 animals) had both young adult (4 male, 1 female) and aged adult animals (n=5 all male). The multiple sessions of rTMS group (total n=5 animals) consisted of young adult animals only (3 males, 2 female).

### Cranial window implantation

Animals were habituated to the animal facility for at least 7 days prior to cranial window implantation surgery. For cranial window imaging, we used a modified protocol from [20]. Briefly, animals were given pre-operative analgesia (buprenorphine, 0.1mg/kg, subcutaneous), anaesthetised with isoflurane and placed in a stereotaxic frame. Local anaesthetic (5mg/kg bupivacaine) was infiltrated under the scalp prior to incision and skin removal. A high-speed dental drill was used to perform a 3mm craniotomy over the right primary motor cortex region (M1) likely blending into the SS1 somatosensory region (window centred over +1mm anterior from bregma, +2.5mm lateral to the midline). Artificial cerebrospinal fluid (ACSF) was applied regularly to cool the bone during drilling. The exposed dura was left intact and was gently cleaned with sterile gelfoam (Pfizer), soaked in ACSF.

Dexamethasone was applied topically to the dura (approximately 20μl of 4mg/ml solution), as this has been shown to enhance window clearance [21]. A 5mm circular glass coverslip was placed over the craniotomy, and the perimeter sealed with Loctite 454 (a low-volatile, low-temperature curing cyanoacrylate gel). A titanium bar with a threaded hole was glued onto the left side, opposite the window. The exposed scalp and skin margin was sealed with dental acrylic (Heraeus). Animals recovered over a 2–3-week period post-surgery before undergoing imaging.

### Multiphoton in vivo imaging

*In vivo* two-photon laser scanning microscopy (2PLSM) was performed using a custom-built upright laser scanning microscope (Scientifica) equipped with galvo-galvo mirrors, a 20X water immersion lens (NA 1.0, Zeiss Plan-Apo) and a high-sensitivity GaAsP non-descanned photomultiplier detector (Hamamatsu). Femtosecond-pulsed infrared excitation was from a mode-locked Ti-sapphire laser tuned to 910nm and equipped with group velocity dispersion compensation (Mai Tai DeepSee, Spectra-Physics). Power delivered to the back aperture was 20-90mW, depending on depth. This setup enabled us to image a good cranial window to >1000µm into the cortex (see Supplementary Figure 1 for point spread function). Images were acquired with ScanImage 3.8.1 software (Vidrio Technologies, LLC) [22] incorporating the Navigator plugin.

Young adult animals were anaesthetised during imaging with an injectable anaesthetic (100mg/kg ketamine, 10mg/kg xylazine, intraperitoneal injection). Anaesthesia was changed to isoflurane for the aged animals due to their variable and prolonged post-anaesthesia recovery times. Isoflurane anaesthesia was delivered with a mask, using concentrations ranging from 1.75-3% isoflurane as required to maintain a stable plane of anaesthesia. This was assessed by regular monitoring of breathing, and isoflurane adjusted to keep respiration within a range of 40-80 breaths per minute. Analgesic state was not assessed (e.g., by pedal withdrawal reflex) as the purpose of the anaesthesia was for sedation. The titanium bar embedded in the acrylic cap during surgery was attached to a customised brass clamp to maintain stability during imaging. Images of dendritic arbours were captured from the upper 100μm of the cortex for the purposes of quantification, and the neuron type (layer 2/3 or layer 5) was identified with deeper imaging. Images were captured at 0.2μm/pixel resolution in the XY plane, with 1μm Z-steps. Dendritic arbours were selected on the basis of clarity and were not restricted to any particular position within the window. Repeat imaging of dendritic segments was carried out using vascular landmarks to locate the regions of interest.

### Subthreshold rTMS

Purpose built rodent specific coils (circular coil 8mm in diameter and height-see [2]) controlled by an arbitrary waveform generator (Agilent 335141B) connected to a bipolar power supply (Kepco BOP 100-4M) were used to administer the rTMS trains. The rTMS pulses were monophasic in shape (400µs rise time, dB/dT of 300T/s) which induced a peak magnetic field of 0.12T at the base of the coil (Hurst GM08 Gaussmeter). At this intensity, the peak sound intensity of the rTMS “clicks” are ∼26dB SPL at the base of the coil [2] which is below the hearing thresholds previously reported for adult C57Bl6J mice [23]. Each stimulation session delivered 600 pulses of intermittent theta-burst stimulation (iTBS) [24] which contained pulse trains consisting of 3 pulses at 50Hz, repeated at 5Hz and delivered for 2s, with an inter-train interval of 8s (total stimulation time 192s).

All imaging and stimulation timepoints were kept consistent across all mice, with rTMS always delivered in the afternoons (range across the entire experiment was from 12:30 – 16:45). rTMS was delivered 3 hours after the end of each imaging session to allow for complete recovery from the anaesthesia, as anaesthesia during the delivery of rTMS has been shown to alter the effect of rTMS in the rodent brain [25, 26]. Stimulation was delivered to lightly restrained mice that were placed in a restraint bag with a breathing hole at one end (Able Scientific). The rTMS coil was positioned over the mouse cranium, such that the coil windings overlaid the cranial window, with the coil offset laterally to minimise direct stimulation of the motor cortex in the other hemisphere (i.e., the motor cortex without a cranial window). During stimulation, the coil was held directly on the restraint bag (∼1mm distance from the base of the coil to the mouse cranium).

### Finite element method modelling of the induced electric field

Finite element method (FEM) modelling was conducted using COMSOL Multiphysics 5.6 (Burlington, NJ, USA) to estimate the magnetic flux density and the electrical field induced in the mouse brain with our rodent specific rTMS coil (see Figure 1). The estimated peak magnetic field was 0.12T at the base of the coil which reduced to 0.07T at the surface of the cortex. The estimated peak electric field at the surface of the cortex was 15.8V/m and as expected, higher electric field values occurred over the right motor cortex and were particularly high near the location of the cranial window due to the offset position of the coil. See Supplementary material for further details on FEM modelling methodology and results.

**Figure 1.**
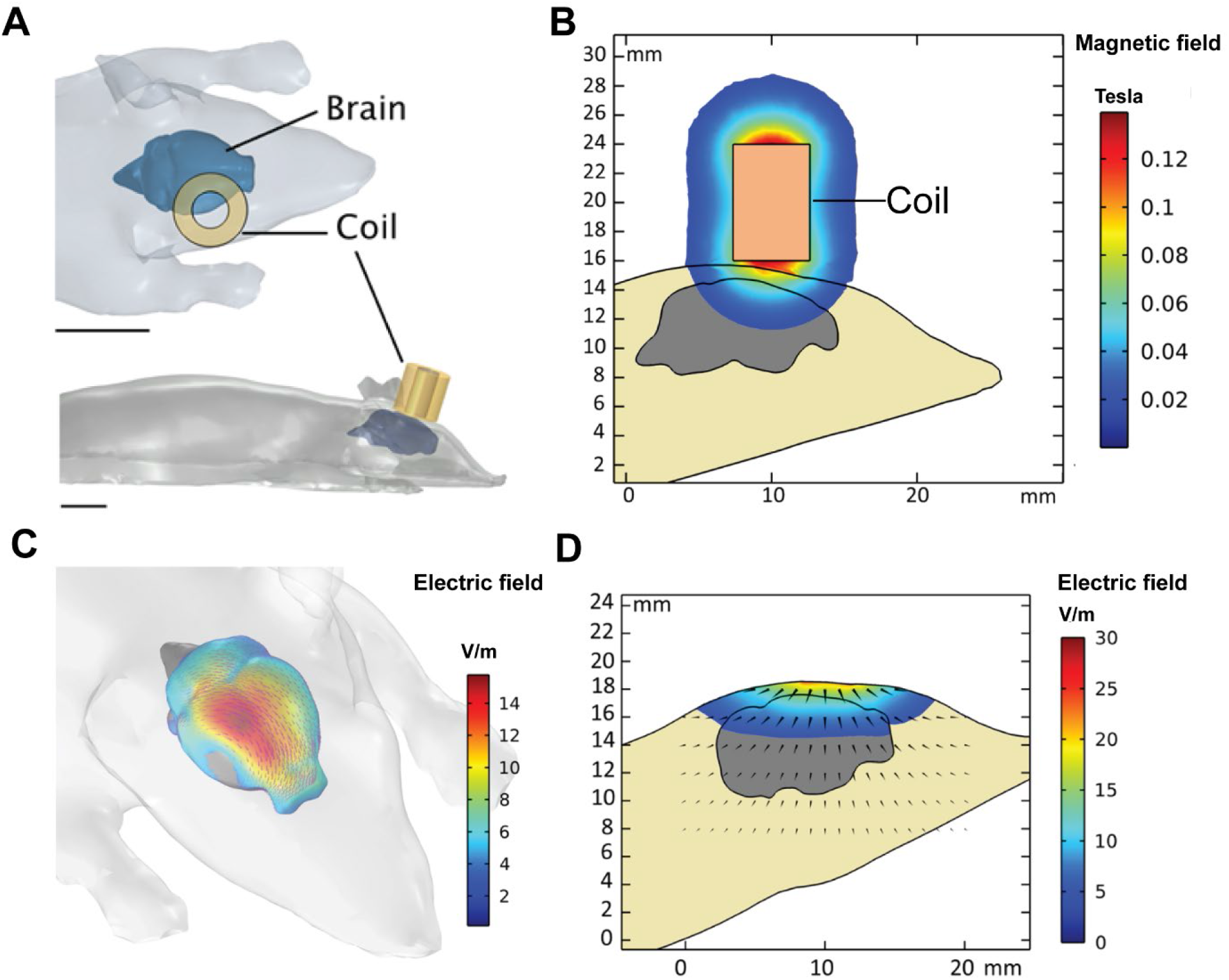
FEM Modelling of the electric field induced over the mouse brain from our rodent-specific TMS coil. (A) Schematic of coil position over the motor cortex/cranial window (scale bars = 1mm). (B) Modelled magnetic field around the coil and through the mouse brain (sagittal plane below the edge of the coil over the motor cortex). (C) Modelled electric field induced across the mouse cortex surface. (D) Modelled electric field induced within the mouse cortex (sagittal plane below the edge of the coil over the motor cortex).

### Imaging timeline

Imaging was separated into 3 time periods (summarised in Figure 2), (i) pre-stimulation, (ii) +21 hours post-stimulation and (iii) +45 hours post-stimulation. For both the single and multiple stimulation groups, pre-stimulation imaging consisted of three imaging sessions conducted over 3 consecutive days separated by 24 hours to capture baseline spine density and dynamics (rate of spine losses and gains). The +21 hours post-stimulation timepoints were conducted 21 hours after each session of rTMS (i.e. once for the single session of LI-rTMS group or a total of four +21 hours post-stimulation timepoints for the multiple stimulation group). A final imaging session was conducted 24 hours after the last +21 hours post-stimulation period and represents the +45 hours post-stimulation timepoint for both the single and multiple stimulation groups.

**Figure 2.**
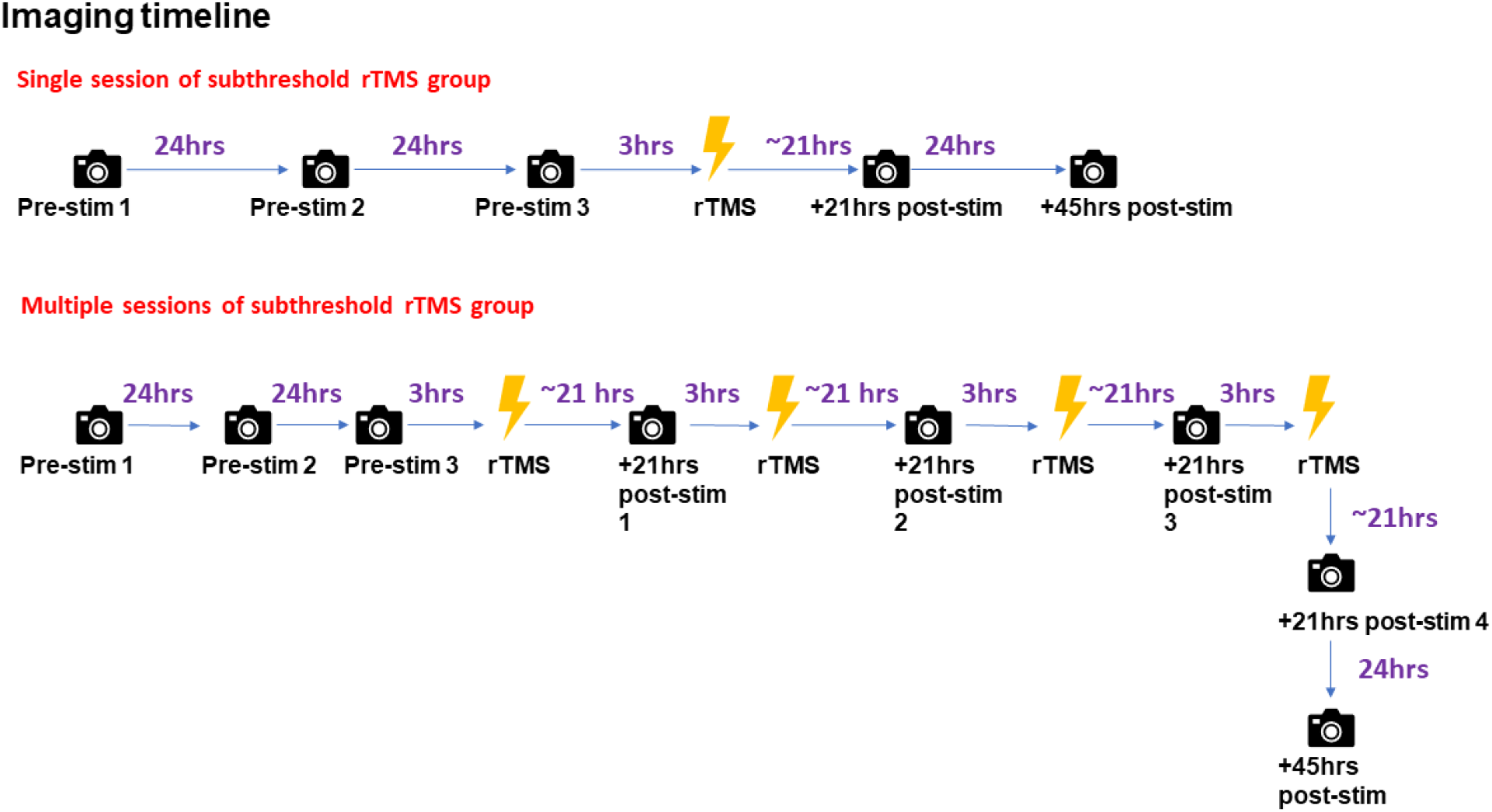
Imaging timeline for the single and multiple sessions of subthreshold rTMS groups.

### Image annotation and spine analysis

Captured stacks were screened for dendrite lengths that were suitable for spine analysis. Although no minimum length was prescribed, only dendritic segments that had at least 11 spines at the final pre-stimulation timepoint were included in the analysis. This resulted in a total of 29 analysable dendritic segments for the single stimulation group (pooled from 10 mice) and 15 analysable dendritic segments for the multiple stimulation group (pooled from 5 mice). Overlapping image stacks were montaged together using XUVTools software [27]. Image stacks were false coloured for depth in Fiji [28] and rotated in 3D to ensure that unique dendritic segments were identified. Where segments joined at a branch point, they were combined and considered to be a single segment. Segments were cropped and then annotated using Matlab scripts included with ScanImage (scim_spineAnalysis.m) to correlate spines across longitudinal time series [20, 22]. Spines were annotated by drawing a line from the centre of the dendritic backbone to the tip of the spine and were only annotated if present across two consecutive z-planes within the stack. Spines were then correlated across the timeseries and exported as spreadsheets of spine lengths where each column was a timepoint and each row corresponded to a single spine across the timeseries. For each dendritic segment, the width of the dendritic backbone was measured at five evenly spaced points along the segment and spine lengths were corrected by subtracting half the mean backbone width.

A custom R script (R core team, 2014) was used to analyse gains and losses, where the criterion for a spine being counted as present was set to 0.4μm (corresponding to two pixels protruding from the edge of the backbone). In addition, a spine had to reach a minimum of 0.6μm (three pixels) at some point along the timeseries, or the entire row was censored. Gains were therefore classified as when a spine went from absent (<0.4μm) to present (≥0.4μm), with loss being the inverse. Transient losses where the spine was absent for only a single timepoint before reappearing at the same location were checked manually. In all except a single case (out of more than 90 apparent transient loss events), these were found to be spurious, and were caused by poor image quality, or rotation of the spine around the backbone to project along the z-axis (detectable by an increase in the brightness of the backbone). All such transient loss events were reclassified as being present throughout. Following data curation and removal of dendritic segments with insufficient spines, the dendritic segments which were analysed statistically tallied (across all timepoints) 17 segments (total length 1807μm) with 2423 annotated spines in the young adult single stimulation group; 12 segments (total length 1691μm) with 1852 annotated spines in the aged adult single stimulation group and 15 segments (total length 1586μm) with 2865 spines annotated in the young adult multiple stimulation group (see Figure 3 and Supplementary Figure 2 for representative images).

**Figure 3.**
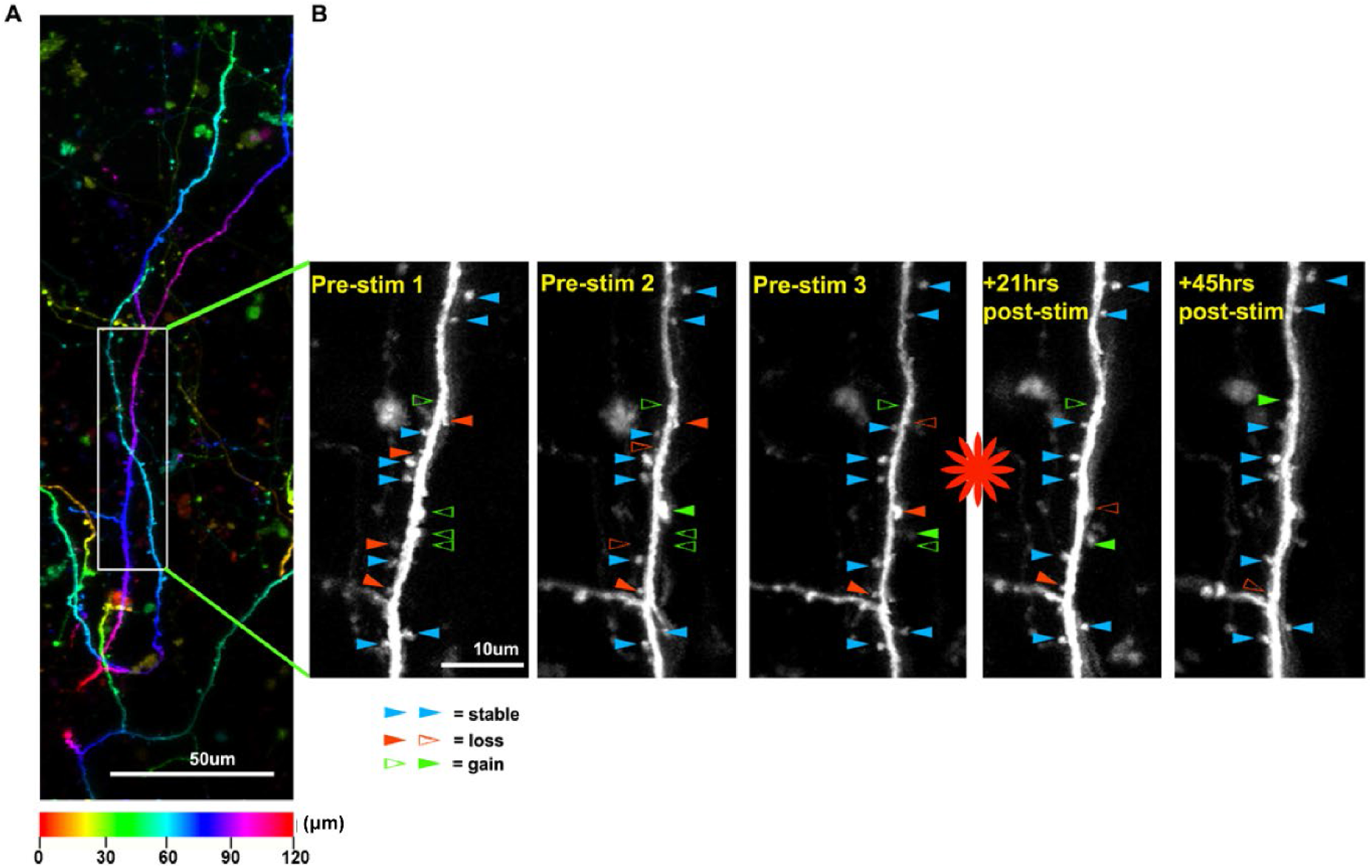
Representative images of an analysed dendritic arbour from the single session of rTMS group. (A) Portion of a dendritic arbour, shown as a maximum projection of a 120μm stack, false colour coded for depth. The colour scale along the bottom shows the correspondence between hue and depth within the stack. Depth of an arbour within the stack does not necessarily correspond to depth from the cortical surface, due to the angle of capture. However, the colours do indicate the relative depths of adjacent or overlapping dendrites. The rectangular box indicates a portion of the annotated subregion shown in panel B). (B) 60μm subregion of a single annotated dendrite from the single stimulation group (total annotated length was 242μm), showing five sequential timepoints at 24hr intervals. The red asterisk indicates subthreshold rTMS stimulation, occurring 3hrs after the Pre-stim 3 image. Spines were annotated in MATLAB and quantitatively analysed using custom R scripts, with the threshold for spine presence set to projection of ≥0.4μm from the edge of the dendritic backbone. Spines over the time series were classified as stable, lost or gained. Blue arrowheads indicate spines present throughout (stable).

Filled red arrowheads indicate spines present that will be lost, with the point of loss shown as a hollow red arrowhead. Hollow green arrowheads indicate nascent spines that have not yet exceeded the threshold, with the solid green arrowhead indicating the point of gain.

### Statistics

For statistical comparison, we compared the 3 time periods (pre-stimulation, +21hrs post-stimulation and +45hrs post-stimulation). Spine density represents the number of spines per µm measured at each imaging session/timepoint. The rate of spine losses and gains represents the number of new spines (per µm) gained or lost (per spine) *since* the last imaging session. The rate of spine losses cannot be calculated for the first pre-stimulation timepoint, as this timepoint provides the baseline counts of spines for each dendritic segment. To facilitate comparison between gains and losses, we similarly excluded this timepoint for the analysis of gains.

Bayesian hierarchical generalized linear regression models were used to estimate (log_e_-) spine density and the rate of spine losses and gains at each 24-hour observation and the average rate (per 24-hour) within the 3 imaging periods (pre-stimulation, +21hrs post-stimulation, +45hrs post stimulation with rTMS) for each stimulation group. Data were counts, therefore a Poisson distribution was assumed, with log-link function. To measure changes in spine density we estimated the expected (log_e-_) number of spines per µm per 24hr. Since turn-over (rate of new spine growth and proportion of spines lost) can vary without changing spine density we estimated the proportion of spines lost per 24hr (*i*.*e*., adjusted for number of spines on the dendrite at the previous 24hr observation) and the rate of spine gain per 24-hours adjusted for dendritic length (log_e-_µm), as longer dendrites are likely to have more spines. Weakly regularizing student-t priors with 3 degrees of freedom (fat tails) were specified, expressing some scepticism that there was any difference in rate of gains or losses between observations or stimulation periods.

Given that the data in the single stimulation and multiple stimulation groups are identical in the way they are collected from the pre-stimulation period up to the first +21hrs post-stimulation observation (see Figure 2), a pooled analysis was run on this combined dataset to supplement our single stimulation analysis.

All statistical analysis was conducted in the R statistical computing environment (R Core Team, 2014). We used the R package ‘brms’ [29] to specify models using Hamiltonian Monte Carlo sampling in Stan [30]. Chain mixing and convergence was assessed using trace plots and R-hat statistics, and model fit was assessed using (graphical) posterior predictive checks. We used Bayesian statistics (see [31] for a review) to determine the level of evidence in favour of the alternative hypothesis (i.e. that subthreshold rTMS had an effect on our dendritic spine measures). Estimated marginal means (averaged over age and/or layer) were computed using the ‘emmeans’ R package [32]. Marginal likelihoods were computed over 27×10^3^ posterior samples (after a 3 ×10^3^ iteration warmup). All code and data to reproduce the analysis will be accessible through online repositories (https://research-repository.uwa.edu.au/en/persons/alex-tang/datasets/ and https://eprints.utas.edu.au/).

We fitted 3-level models with random intercepts for dendritic segments nested within animal, with age and cell body location (layers 2/3 or 5) included as fixed effects. We report rate of gains per day, per micron, and rate of losses per day, per spine (lagged to the previous observation as described above). Uncertainty is expressed as 95% (posterior) credible intervals (CI). As we used an internal control design, our analysis (including hypothesis testing) estimates the dendritic spine dynamics at the population-level at the pre-, +21hrs post-stimulation and +45hrs post-stimulation time periods. Bayes Factors (BF-evidence ratios) are reported to aid in the interpretation of our results and we used the following BF cut-offs as evidence in favour of the alternative hypothesis [33]. Only results that had “strong” or “extreme evidence” (i.e. a BF≥10) were interpreted to have biological relevance (see Table 1).

**Table 1.** Summary of results after subthreshold rTMS (i.e., post-stimulation time points compared to pre-stimulation). The direction of change (increased/decreased) is indicated only for results with a BF≥10 (i.e., with ≥strong evidence of a change). NOTE: “Age” is not included in the table as it was investigated in the single stimulation group only and did not show strong evidence for any spine measure. Pooled analysis encompasses +21hrs post-stimulation time points only.

|  | Dendritic spine density |  | Rate of dendritic spine losses |  | Rate of dendritic spine gains |  |
| --- | --- | --- | --- | --- | --- | --- |
|  | +21 hours post-stim | +45 hours post-stim | +21 hours post-stim | +45 hours post-stim | +21 hours post-stim | +45 hours post-stim |
| Single session of subthreshold rTMS | - | ↓ | ↑ | - | - | ↓ |
|  | BF=1.8 | BF=18.2 | BF=13.4 | BF=1.1 | BF=2.0 | BF=564.4 |
| Multiple sessions of subthreshold rTMS | - | - | ↑ | - | - | - |
|  | BF=1.7 | BF=1.5 | BF=20.4 | BF=1.6 | BF=1.9 | BF=3.5 |
| Pooled analysis up to first +21hrs post-stimulation timepoint | - |  | ↑ |  | ↑ |  |
|  | BF=2.1 |  | BF=34.9 |  | BF=35.0 |  |

- **BF = 1** - No evidence
- **1 < BF <= 3** - Anecdotal
- **3 < BF <= 10** - Moderate
- **10 < BF <= 30** - Strong
- **BF>100** - Extreme

While our model adjusted each spine measure for cell layer, the majority our data consists of layer 5 neurons (15/20 neurons for the single stimulation group and 10/11 neurons for the multiple stimulation group). Therefore, no evidence ratios are reported for the comparison between layer 2/3 and 5 as we are unable to make meaningful interpretations from an unbalanced sample.

## Results

### Single session of subthreshold rTMS (young adult and aged mice)

#### Density

Compared to pre-stimulation (0.216 spines μm^-1^, 95%CI 0.149, 0.284, averaged over layer and age, n=29 dendritic segments from 20 neurons from 10 mice) there was anecdotal evidence for a decrease in spine density at +21hrs post-stimulation (0.213 spines μm^-1^, 95%CI 0.146, 0.282, BF=1.78). In contrast, there was strong evidence for a decrease in the mean dendritic spine density at +45hrs post-stimulation (0.201 spines μm^-1^, 95%CI 0.137, 0.265, BF =18.2, see Figure 4A). Further analysis showed anecdotal evidence for a greater effect of subthreshold rTMS on dendritic spine density in young adults (n=5) compared to aged animals (n=5) (BF=1.5).

**Figure 4:**
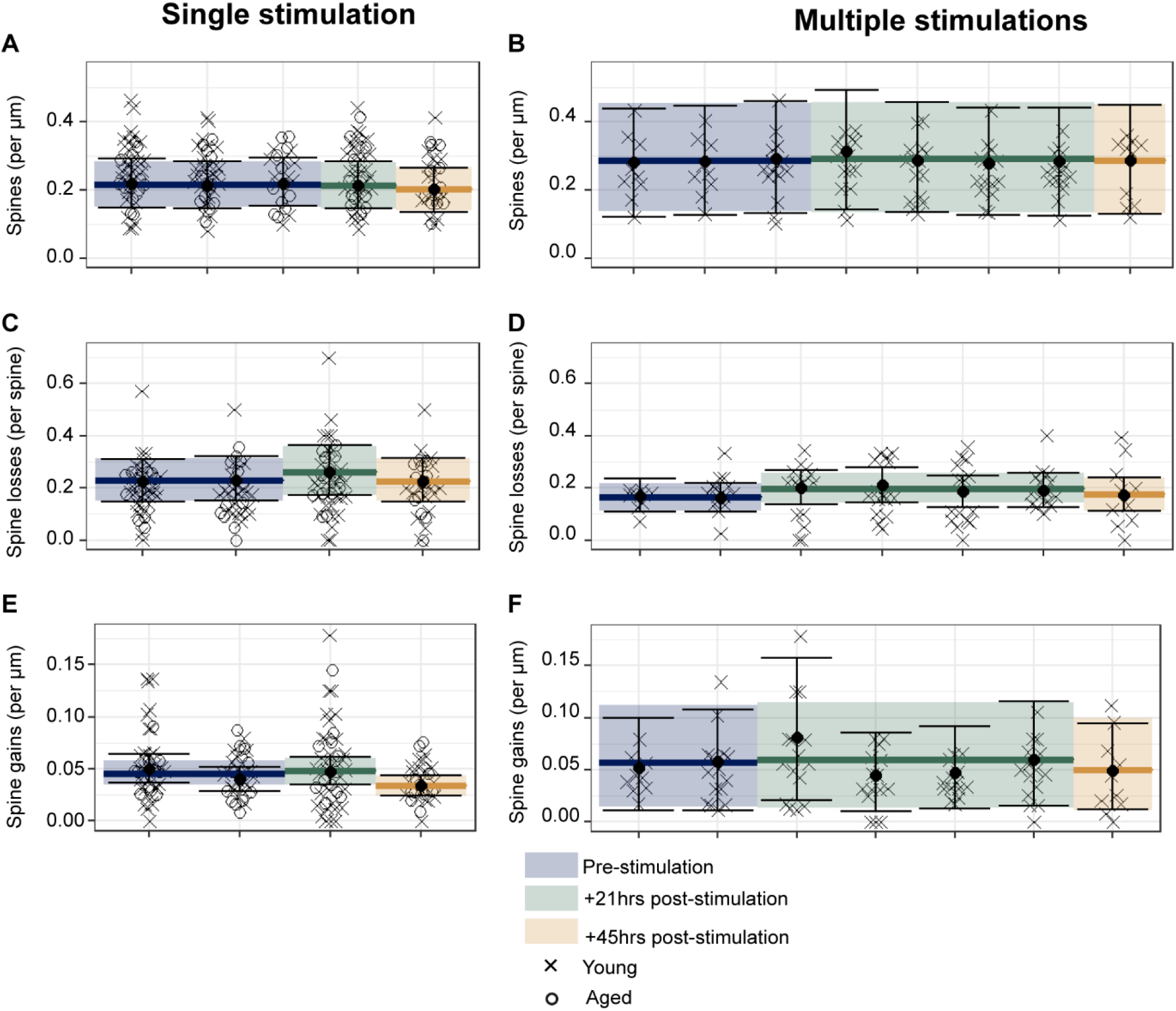
Single (left side) and multiple sessions of subthreshold rTMS (right side) drive structural synaptic plasticity in pyramidal neurons of the motor cortex. (A and B) Analysis of dendritic spine density showed strong evidence for a decrease in the mean dendritic spine density +45hrs post a single session of subthreshold rTMS, compared to pre-stimulation. (C and D) Analysis of the rate of dendritic spine losses showed strong evidence for an increase in the mean rate of spine losses +21hrs post-stimulation in both the single and multiple sessions of subthreshold rTMS groups compared to pre-stimulation. (E and F) Analysis of the rate of dendritic spine gains showed strong evidence for a decrease in the mean rate of dendritic spine gains +45hrs post a single session of subthreshold rTMS compared to pre-stimulation. Data are shown as the aggregate means (solid-coloured lines) for each time period (pre-stimulation=blue, +21hrs post-stimulation=green, +45hrs post-stimulation=orange) overlaid on the mean (•) at each imaging observation. Error bars represent the 95% **credible intervals** for each timepoint, whereas the shaded boxes represent the 95% credible intervals for each time period. Each data point represents data from an individual dendritic segment with data from young (x) and aged (o) animals.

#### Rate of spine loss

Compared to pre-stimulation (0.228 spine^-1^, 95%CI 0.156, 0.315, averaged over layer and age), there was strong evidence for an increase in the mean rate of spine losses following subthreshold rTMS at +21hrs post-stimulation (0.262 spine^-1^, 95%CI 0.173, 0.361, BF =13.4) but not at +45hrs post-stimulation (0.227 spine^-1^, 95%CI 0.151, 0.313, BF=1.1, see Figure 4C). In addition, there was strong evidence for a greater rate of spine losses at +21 and compared to +45hrs post-stimulation (BF=10.3), further suggesting that there was an increase in the rate of spine loss following subthreshold rTMS that resolves by +45hrs post-stimulation. Analysis of age showed moderate evidence for a greater increase in the rate of spine losses in young adult animals compared to aged adult animals (BF=4.2).

#### Rate of spine gain

Compared to pre-stimulation (0.046 spines μm^-1^, 95%CI 0.035, 0.058, averaged over layer and age), there was anecdotal evidence of an increase in the rate of spine gain following subthreshold rTMS +21hrs post-stimulation (0.048 spines μm^-1^, 95%CI 0.035, 0.06, averaged over layer and age, BF = 2.0, see Figure 4E). In contrast, there was extreme evidence for a decrease in the rate of spine gain at +45hrs post-stimulation (0.034 spines μm^-1^, 95%CI 0.025, 0.044, averaged over layer and age, pre-stimulation> +45hrs post-stimulation: BF = 564.4, +45hrs post-stimulation< +21hrs post-stimulation: BF = 661.6). There was anecdotal evidence for a greater change in the rate of spine gain between young adult and aged animals (BF=0.9).

### Multiple sessions of subthreshold rTMS (young adult mice)

#### Density

Compared to the pre-stimulation period (0.289 spines μm^-1^, 95%CI 0.135, 0.462, n=15 dendritic segments from 11 neurons from 5 mice, averaged over layer), there was anecdotal evidence for an increase in the mean density of dendritic spines following subthreshold rTMS at the +21hrs post-stimulation timepoints (averaged over the 4 +21hrs post-stimulation timepoints, 0.293 spines μm^-1^, 95%CI 0.141, 0.473, BF = 1.7, see Figure 4B) and a decrease at +45hrs post-stimulation (0.287 spines μm^-1^, 95%CI 0.137, 0.67, BF = 1.5). In addition, there was anecdotal evidence for a decrease in density between the +21hrs post-stimulation timepoints and +45hrs post-stimulation (BF=1.62).

#### Rate of spine loss

Compared to pre-stimulation (0.166 spine^-1^, 95%CI 0.116, 0.219, averaged over layer), there was strong evidence for an increase in the mean rate of spine losses at the +21hrs post-stimulation timepoints (averaged over the 4 +21hrs post-stimulation timepoints 0.199 spine^-1^, 95%CI 0.146, 0.255, BF= 20.4, see Figure 4D) but not at +45hrs post-stimulation (0.175 spine^-1^, 95%CI 0.111, 0.240, BF=1.6).

#### Rate of spine gain

Compared to pre-stimulation (0.057 spines μm^-1^, 95%CI 0.014, 0.109, averaged over layer), there was anecdotal evidence for a decrease in the mean rate of spine gain at the +21hrs post-stimulation timepoints (averaged over the 4 +21hrs post-stimulation timepoints +0.059 spines μm^-1^, 95%CI 0.016, 0.115, BF= 1.9, see Figure 4F). Following subthreshold rTMS, there was moderate evidence for a decrease in the rate of spine gain at +45hrs post-stimulation (0.050 spines μm^-1^ 95%CI 0.013, 0.098, BF = 3.5).

### Pooled analysis of single and multiple stimulation data (from pre-stimulation to the first +21hrs post-stimulation timepoint)

As the single and multiple stimulation groups are identical experimentally, to the first +21hrs post-stimulation imaging timepoint (i.e., up to +21hrs post a single stimulation), we pooled the data from both groups up until this point to supplement our single stimulation analysis.

#### Density

Increasing the sample size in the pooled analysis (total n=44 dendritic segments from 26 arbours, from 15 mice) showed anecdotal evidence for an increase in the mean dendritic spine density +21hrs post a single session of subthreshold rTMS (0.229 spines μm^-1^, 95%CI 0.173, 0.288) compared to pre-stimulation (0.223 spines μm^-1^, 95%CI 0.171, 0.284, BF = 2.1, see Supplementary Figure 3A).

#### Rate of spine loss

Pooled analysis showed strong evidence of an increase in the rate of spine losses at +21hrs post a single session of subthreshold rTMS (0.238 spine^-1^, 95% CI 0.150, 0.263) compared to pre-stimulation (0.204 spine^-1^, 95% CI 0.174, 0.307, BF=34.9, see Supplementary Figure 3B).

#### Rate of spine gains

Pooled analysis showed strong evidence (BF=35.0, see Supplementary Figure 3C) for an increase in the rate of spine gain at the +21hrs post a single session of subthreshold rTMS (0.053 spines μm^-1^ 95%CI 0.035, 0.057) compared to pre-stimulation (0.045 spines μm^-1^ 95%CI 0.039, 0.066).Given that this result was not evident in the single stimulation analysis, this suggests that this effect is not as robust as the change seen in spine losses, as it could only be detected by increasing the sample size. Further analysis of the pooled data did not indicate that animals in the multiple stimulation group had a greater increase in gains during the first +21hrs post-stimulation period compared to the animals in the single stimulation group (see Supplementary Material).

## Discussion

To our knowledge, this is the first *in vivo* study to provide evidence that subthreshold rTMS, drives structural synaptic plasticity in the motor cortex. Following a single session of subthreshold rTMS, there was an increase to the rate of dendritic spine losses (present at +21hrs post-stimulation) and a long-lasting decrease in dendritic spine density and to the rate of dendritic spine formation (present at +45hrs post-stimulation) which did not differ between young and aged animals (see Figure 4). Surprisingly, multiple days/sessions of subthreshold rTMS did not have a cumulative effect or longer lasting effect on structural plasticity compared to a single session of subthreshold rTMS but instead maintained the more acute effects induced by a single session (increased rate of spine losses at +21hrs post-stimulation) which returned to baseline values by +45hrs after the last subthreshold rTMS session.

Given that previous rTMS studies in humans using greater intensities have shown age-related differences in rTMS-induced plasticity in other measures of neural plasticity (motor evoked potentials and the default mode network) with the iTBS protocol [12, 13], we might have expected similar results in our study. However, we failed to find strong evidence of a difference between young adult and aged animals in the structural plasticity of dendritic spines induced by a single session of subthreshold rTMS. This may be explained by the absence of age-dependent differences in dendritic spine properties pre-stimulation, such that subthreshold rTMS modulated both the young and aged brain as they did not differ in the amounts of synaptic connectivity and remodelling capacity prior to stimulation. Interestingly, these results are in direct contrast to the work of Davidson et al., who reported age-dependent differences in pyramidal neuron dendritic spine density and dynamics in the forelimb region of the motor cortex in Thy1-GFP mice [10]. Specifically, in both our young and aged animals, the mean dendritic spine density and rate of spine *gains* in the pre-stimulation period was approximately half of that reported by Davidson et al., whereas the rate of spine *losses* from our young adults was almost twice as large. In addition to several methodological differences that may explain these contrasting findings, a key point of difference between our studies was that our cranial windows were centred more posterior relative to bregma to that used in Davidson et al. Therefore, while we did not find age-dependent effects on baseline or subthreshold rTMS induced spine plasticity in the area of the motor cortex we imaged, it’s possible that such differences do occur, depending on the anatomical region imaged or circuit stimulated.

In a recent study, Cambiaghi et al., found that 5 daily sessions of 15Hz rTMS (intensity not reported), increased the spine density on apical and basal dendrites of layer 2/3 pyramidal neurons in the motor cortex of young adult mice (2.5-3 month old) when measured 24 hours post-stimulation [34]. The use of Golgi-staining in brain slices (i.e., investigated post-mortem) suggested that the rTMS protocol used, preferentially increased the density of “thin” spines in apical dendrites, which are suggested to be “less stable” and potentially in a state of “learning” [35]. Measuring similar outcomes in our data is difficult, especially across several timepoints as quantifying spine morphologies from *in vivo* images presents several challenges (e.g., intrinsic movement of the spines between imaging sessions). However, a qualitative analysis of a small subset of single stimulation data (see Supplementary material – Spine morphologies and Supplementary Tables 1 and 2) did not show an obvious effect of subthreshold rTMS to the rates of losses or gains towards a specific spine subtype. However, given the effect sizes observed, a much larger sample size would be needed to perform a meaningful statistical comparison. We suggest this would be an interesting experiment to perform in the future, ideally quantified post-mortem, where spine morphologies can be captured with greater resolution (e.g., with confocal microscopy and objectives with high numerical apertures).

In young adult mice, we have previously shown that 10 daily sessions of subthreshold rTMS (with the same 600 pulses of iTBS protocol) to the motor cortex paired with skilled motor training improves motor behaviour and the rate of learning [3]. In that study, our behavioural analysis suggested that multiple sessions of subthreshold rTMS maintained the transient effects induced by a single stimulation. Similarly, repeated blocks of suprathreshold iTBS within a single stimulation session to the rat barrel cortex was shown to have a cumulative effect on cortical activity [36]. Interestingly, multiple sessions of subthreshold rTMS maintained the initial structural plasticity induced by a single session of rTMS, with both groups showing an increase in the rate spine losses at the +21hrs post-stimulation timepoint (pooled analysis-see Supplementary Figure 3). However, the effects of multiple sessions were shorter lasting (+21hrs post-stimulation) compared to a single session which also showed decreased spine density and rate of spine gains at the +45hrs post-stimulation timepoint. While it is unclear why a single-session of subthreshold rTMS had longer lasting effects on structural plasticity than multiple-sessions, the net changes observed with both groups suggest a common outcome of refinement in synaptic connectivity as an underlying mechanism of subthreshold rTMS-induced plasticity.

This is the first demonstration, to our knowledge, that subthreshold rTMS has delayed effects on neural plasticity, as our previous work has shown that subthreshold rTMS (using the iTBS protocol) drives acute changes to motor behaviour [3] and neuronal excitability [1]. While further investigation is needed to determine the functional consequences of structural plasticity induced with subthreshold rTMS, we speculate that the refinement in synaptic connectivity induced may promote neural plasticity, by removing or preventing redundant synaptic connections, thereby producing a more efficient neural circuit. This is consistent with the use of low intensity rTMS (0.012T) in ephrin-A2A5-/- mice to reorganise abnormal circuitry by removing aberrant connections [37, 38].This delayed form of subthreshold rTMS induced structural plasticity in the motor cortex may work in tandem with, or be a result of the immediate effects of subthreshold rTMS, such that the acute changes in neuronal excitability enhance motor behaviour which is later supported by the delayed refinement/pruning of the neural circuitry. However, future studies combining subthreshold rTMS with motor training and *in vivo* imaging is needed to confirm our theory and we do not rule out the possibility that the changes observed independent of a task, may be random and may not target task-relevant connectivity.

The use of rodent-specific coils offered us the advantage of delivering focal stimulation (e.g. restricting stimulation to one motor cortex) but at the expense of stimulation intensity [2]. Therefore, while we have shown that subthreshold rTMS induces synaptic plasticity, it remains unknown what effect suprathreshold rTMS has on structural synaptic plasticity *in vivo*. Investigating such changes in a mouse model is technically challenging as “small” commercial coils (e.g. 25mm figure of 8 coils) stimulate the entire brain [39]. In addition, despite promising improvements in rodent-specific TMS coils [40], there are currently no rTMS coils appropriately sized for the mouse brain that can deliver high intensity rTMS, due to the thermal stress induced with each pulse [2, 41], particularly at high stimulation frequencies such as iTBS. As a result, characterising structural synaptic plasticity with *in vivo* imaging following high intensity rTMS may be more practical in a rat model, where commercial coils can be used with some degree of focality [26, 42, 43].

Despite the extremely low sound generation (<26dBL sound pressure) produced by our rodent coils during rTMS, a limitation of our study is the absence of a sham stimulation group. While internal controls/ comparisons to pre-stimulation measures are commonly used in human rTMS studies (e.g., motor evoked potential properties normalised to pre-stimulation values), we cannot rule out the possibility that our results were influenced by factors outside of subthreshold rTMS such as daily anaesthesia and/or stress during the light restraint used to deliver rTMS. However, given that we did not observe a consistent change over time (e.g., a change in dendritic spine density that increased over time) or consistent changes between the stimulation groups (e.g., a change in a dendritic spine property that was seen at the 5^th^ imaging session in both the single and multiple stimulation groups), repeated anaesthesia is unlikely to have been a significant factor. Furthermore, we analysed a subset of data (n=5 dendritic segments from 3 young adult mice) taken from a similar motor cortex imaging study that was being completed in the lab. This supplementary data did not show strong evidence for a difference between baseline and post-restraint (see Supplementary Material-Effect of restraint on spine measures). While these observations do not replace a sham group, they do provide some data to suggest that the repeated anaesthesia and restraint used in our study are unlikely to have caused the structural synaptic changes we observed after subthreshold rTMS.

In conclusion, our results show that subthreshold rTMS drives structural synaptic plasticity in the mouse motor cortex. These findings uncover structural synaptic plasticity as a key mechanism of subthreshold rTMS induced neural plasticity and more broadly, highlights the ability of rTMS to alter synaptic connectivity in the brain.

## Acknowledgements

The authors thank Claire Hadrill for her assistance with the initial spine annotations and Dr Saied Mehrkanoon for assistance with the modification and re-generation of the MATLAB scripts used for spine annotations. This project was funded by the National Health and Medical Research Council (NHMRC - APP1050261). JR was supported by research fellowships from the NHMRC (APP1002258) and Multiple Sclerosis Western Australia. MRH was supported by research fellowships from the Australian Research Council (DE120100729 and FT150100406). AT was supported by doctoral scholarship from the Australian Government, the Bruce and Betty Green Foundation and post-doctoral funding from the Raine Medical Research Foundation (RPG06-20).

