## Supplemental Material for "Subthreshold repetitive transcranial magnetic stimulation drives structural synaptic plasticity in the young and aged motor cortex"

### Supplementary Material

### FEM modelling methodology

A 3D FEM mouse model was created from the anatomical atlas of the adult male nude mouse “Digimouse” [1], which was derived from cryosection images and 0.1mm isotropic X-ray CT images. 3D surfaces were segmented out using ITK-SNAP [2] based on CT image signal intensity values and were imported into COMSOL for optimisation and meshing. The “Digimouse” model brain was extracted out and replaced by higher resolution MRI images of the P56 mouse atlas brain [3] using the FMRIB software library [4] Brain Extraction Tool (BET), the P56 brain was scaled and mapped to the original brain space and the “Digimouse” spinal cord was kept intact.

The final mouse and coil model consisted of ~2 million domain elements including 726,923 tetrahedra, the mouse body measured 89.75 mm long with a volume of 238.24 $cm^2$. The median thickness between the scalp and cortex surfaces below the coil was 1.05 mm The mouse brain measured 14.13 mm long with a volume of 0.38$cm^2$, which is the reported average volume for a mouse [5] and due to mouse brain volume being stable at three weeks of age onwards and especially from eight week of age this model is valid for both the “young adult” and “aged” cohorts from this current study [6].

The coil model consisted of a homogenised multi-turn coil with 780 turns, an outer diameter of 8 mm, an inner diameter of 4 mm, a height of 8 mm, a wire diameter of 0.125 mm and a soft iron core, as described by Tang et al. [7]. The coil was driven by a 100 V input in the frequency domain at 2.5 kHz to represent the 400 μs biphasic pulse rise time of the coil input.

The coil was placed 1mm above the mouse skull over the right brain hemisphere with the edge of the coil windings over the centre of the implanted cranial window, 1mm anterior from Bregma and 2.5mm lateral to the midline, this represents the experimental placement of the coil on the mouse models during the rTMS sessions. Current was run through the coil in a direction that induced an anterior to posterior current across the superior surface of the right hemisphere in the mouse brain model.

Default isotropic conductivities were set for each layer, which included soft tissues (σ = 0.465S/m), skull (σ = 0.02S/m), CSF (σ = 1.654S/m), grey matter (σ = 0.106/m) and white matter (σ = 0.126S/m), other dielectric properties were set to values used in previous studies of TMS in rodents [8-11], these values are taken from the Foundation for Research on Information Technologies in Society dielectric tissue properties database [12]. The rTMS induced electrical field ($\mathbf{E}$) and magnetic flux density (**B**) were calculated using COMSOL's Magnetic and Electric Fields equations, which solves Maxwell's equations by finding the magnetic vector potential field ($\mathbf{A}$) in the frequency domain, where the rate of change in coil voltage determined the electrical field induced in the brain, given by

$$\mathbf{E} = -\nabla \mathrm{V} - \mathrm{j}\omega\mathbf{A}$$

Where $\nabla$ is the curl, $\omega$ is the frequency (2.5 kHz) and $\mathrm{j}$ is the free current density. Similarly, the magnetic flux density is solved by

$$\mathbf{B} = \nabla \times \mathbf{A}$$

Modelling outputs were calculated using COMSOL's magnetic and electric fields (mef) physics interface with the surrounding boundary containing an infinite domain, which allows the model to remain unaffected by boundary conditions. All model domains were given meshes with at the minimum, an "extra fine" element size and the coil and its core were meshed using the swept function similar the modelling conducted by Tang et al. for the rat brain [7]. While the dielectric properties of the air, copper and iron in the model are unaffected by the frequency of electrical stimulation it is important to note biological tissues have different frequency dependent dielectric properties that alter the resulting E-field induced in the brain [13, 14].

**FEM modelling results**

FEM modelling of our rodent-specific coil estimated the peak magnetic field generated to be 0.12T directly below the base of the coil windings. The magnetic field reduced to just below 0.12T at the scalp surface and ~0.70T at the surface of the cortex, it then reduced further to ~40mT at a depth of 2 mm (see Figure 1).

The peak induced electric field in the mouse scalp was 30V/m which reduced to 15.8V/m at the surface of the cortex. At a depth of 1mm the electric field was ~10.2V/m and ~5V/m at 2mm deep. As expected, higher electric field values were measured over the right hemisphere and were particularly high at the location of the implanted cranial window.

**Point spread function of the microscope**

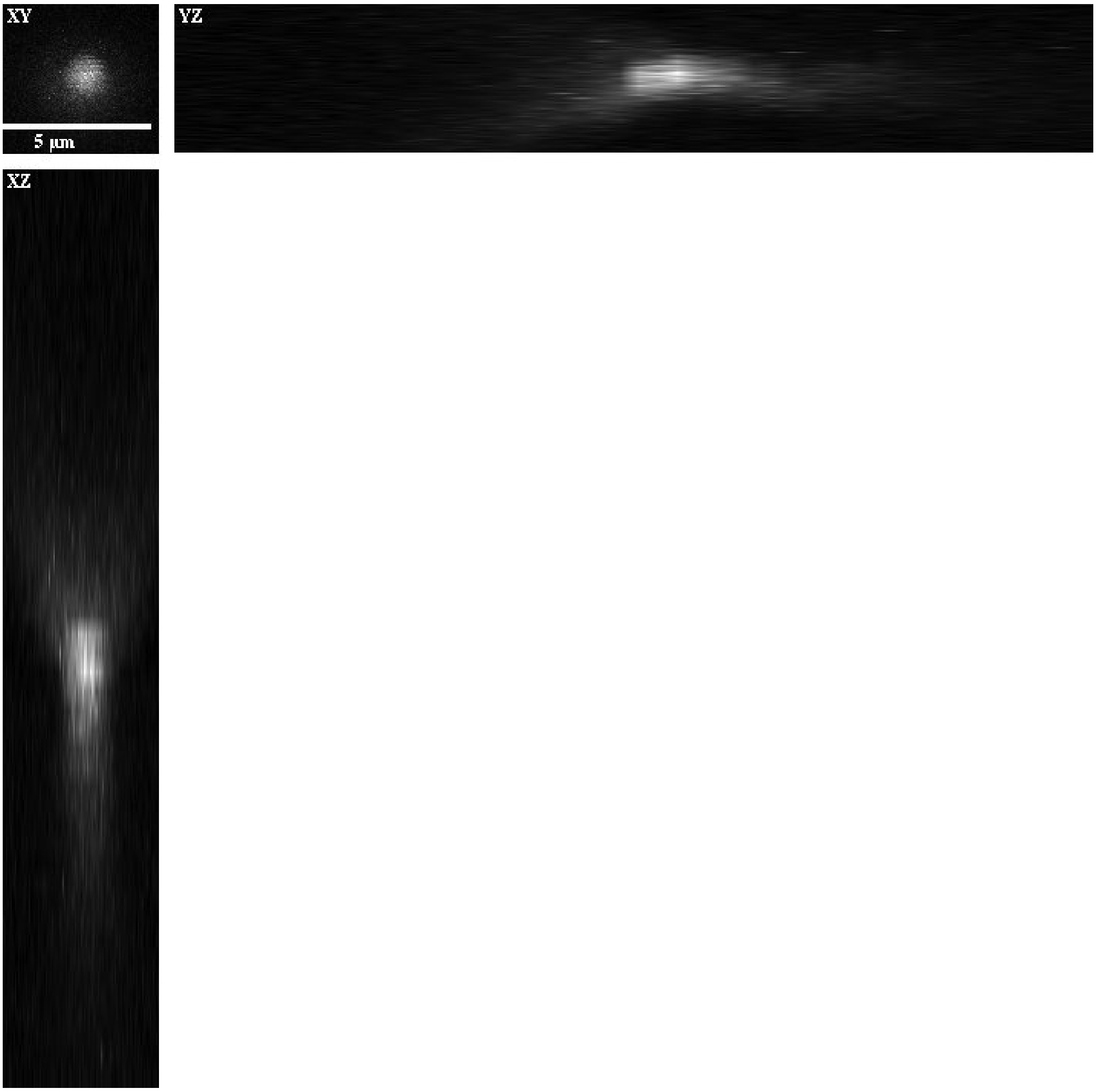

**Supplementary Figure 1. Point spread function of the 2-photon microscope used for imaging. Data was generated using 175nm green fluorescent beads. Full width at half maximum values of the point spread function were 0.692µm (x axis), 0.765µm (y axis) and 1.54µm (z axis).**

**Raw images of multiple stimulation data**

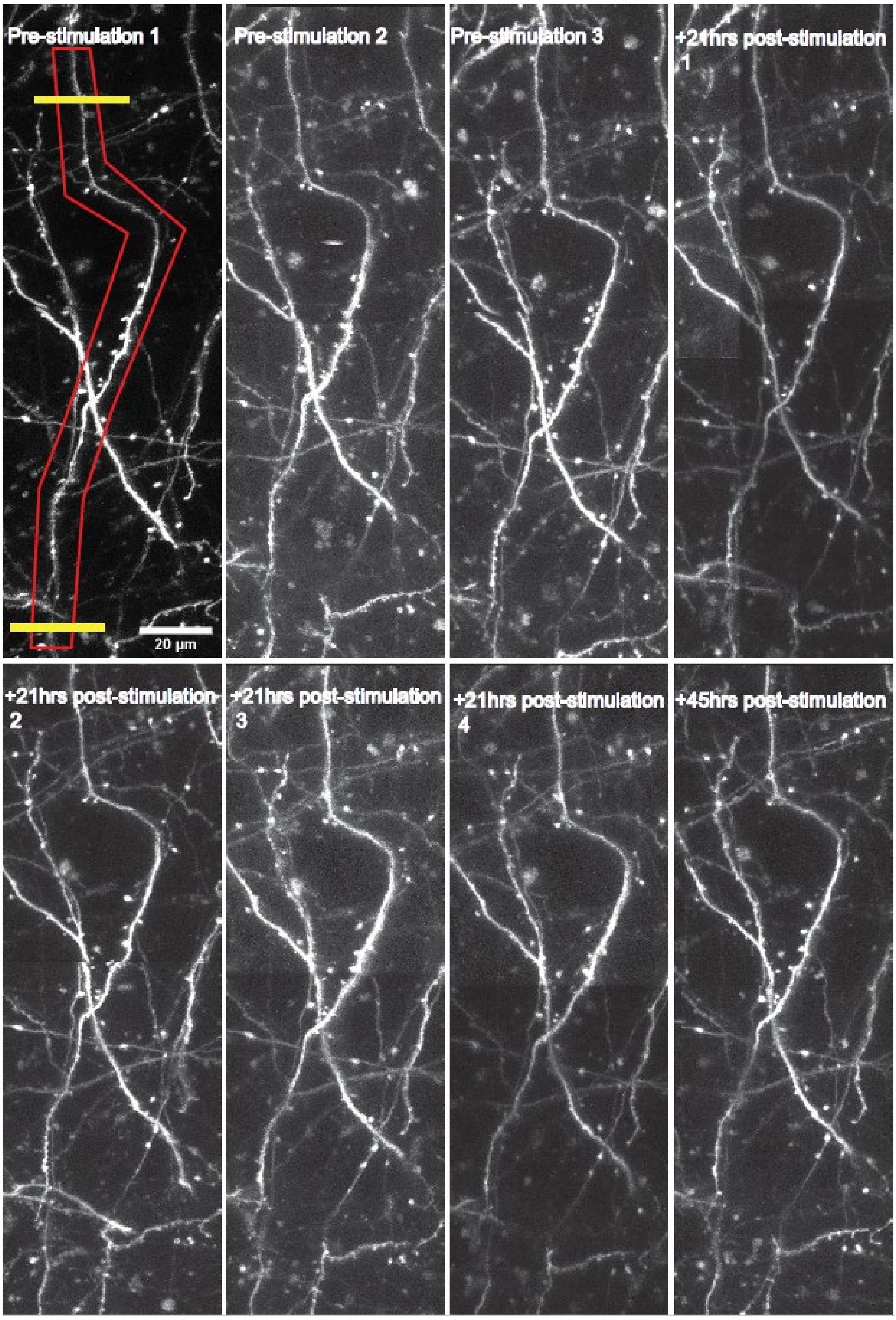

**Supplementary Figure 2. Raw representative images from the multiple stimulation group used for analysis.** Representative images of a dendritic segment (outlined by the red box) analysed for the multiple stimulation group. Spines situated between the two yellow lines were analysed at each time point for this dendritic segment.

**Pooled analysis**

Given that the pooled analysis showed strong evidence for a change to the rate of spine gains at +21hrs post a single stimulation, that was not evident in the single or multiple stimulation analysis alone, further analysis was conducted to determine whether this result was being driven by data from a particular stimulation group (single or multiple stimulation). We ran 3 models, Model 1 with an interaction term between stimulation group and imaging timepoints, Model 2 with stimulation group as a main effect and Model 3 that does not account for any effect of stimulation. Comparison between Models 1 and 2 did not show strong evidence for an interaction effect (BF=1.05), suggesting no difference in the change of the rate of dendritic spine over the imaging timepoints between the single and multiple stimulation groups. Similarly, a comparison between Models 2 and 3 did not show strong evidence for a difference between stimulation groups (BF=0.19), suggesting no difference in the data-generating process between the two stimulation groups.

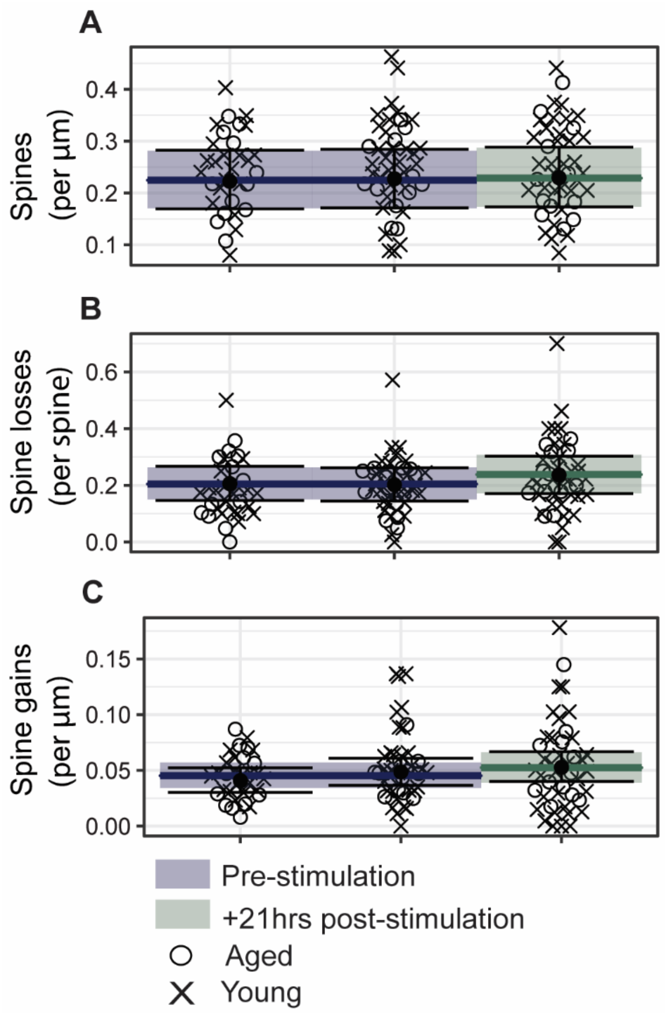

**Supplementary Figure 3. Pooled analysis further suggests that a single session of subthreshold rTMS drives structural synaptic plasticity in the motor cortex.**

(A) Pooled analysis of dendritic spine density did not show strong evidence for a change in density post a single stimulation.

(B) Pooled analysis of dendritic spine losses shows strong evidence for an increase in the rate of spine losses +21hrs post a single stimulation.

(C) Pooled analysis of dendritic spine gains shows strong evidence for an increase in the rate of spine gains +21hrs post a single stimulation.

Data are shown as the aggregate means (solid-coloured lines) for each time period (pre-stimulation=blue, +21hrs post-stimulation=green) alongside the mean (•) at each imaging observation. Error bars represent the 95% **credible intervals** for each individual time point, whereas the shaded boxes represent the average 95% credible intervals for each time period. Each data point represents data from an individual dendritic arbour with data from young (x) and aged (o) animals.

**Spine morphologies**

Longitudinal quantification of spine morphology from *in vivo* images presents several challenges, however we analysed the spine morphologies in a subset of data from the single stimulation group (n=7 segments from 3 mice). As expected, there was substantial variability in the proportion of each spine subtype present on the dendrites, both within and across animals (see Supplementary table 1). Across all the segments we counted 277 spines, with the distribution being 105 mushroom (37%), 131 stubby (48%) and 41 thin (15%). Qualitative analysis of this small subset of data did not show an obvious effect of subthreshold rTMS on the rate of gains or losses on a specific spine subtype at any of the time points. However, given the small effects on subthreshold rTMS on these measures, a much larger sample size (neurons and dendritic segments) would be needed to perform a meaningful statistical comparison.

**Supplementary Table 1. Overview of the proportion of spine subtypes from a subset of the single stimulation data (n= 7 segments from 3 mice).**

| Animal and dendritic segment ID | Animal Age | Dendritic length (µm) | Number of mushroom spines present | Number of stubby spines present | Number of thin spines present |
|---|---|---|---|---|---|
| Animal 223 neuron 001, segment 2 | Aged | 214 | 14 | 11 | 2 |
| Animal 223, neuron 002, segment 001 | Aged | 143 | 10 | 24 | 6 |

| | | | | | |
|---|---|---|---|---|---|
| Animal 224, neuron 001, segment 001 | Aged | 138 | 17 | 34 | 0 |
| Animal 224, neuron 001, segment 002 | Aged | 172 | 20 | 20 | 20 |
| Animal 224 neuron 002, segment 001 | Aged | 242 | 19 | 17 | 3 |
| Animal 129, neuron 001, segment 001 | Young | 152 | 18 | 15 | 5 |
| Animal 129, neuron 002, segment 001 | Young | 225 | 7 | 10 | 21 |
| Subtotal | | 1286 | 105 | 131 | 41 |

**Supplementary Table 2. Overview of the number of gains and losses of each spine subtype from the combined data subset shown in Supplementary table 1. Spine gains and losses at each of the imaging timepoints are shown.**

| **Spine subtype** | **Gains Pre-stim 1** | **Gains +21hrs post-stim** | **Gains +45hrs post-stim** | **Losses Pre-stim 1** | **Losses +21hrs post-stim** | **Losses+45hrs post-stim** |
|---|---|---|---|---|---|---|
| Mushroom | 4 | 0 | 1 | 5 | 0 | 0 |
| Stubby | 15 | 6 | 9 | 6 | 13 | 1 |
| Thin | 4 | 5 | 5 | 3 | 4 | 4 |

**Effect of restraint on spine measures**

This study used an internal control design (i.e., comparing spine changes pre-and post- rTMS). A limitation of this design is the potential for other factors such as stress from the restraint to affect our results. To estimate the effect of restraint alone on structural plasticity, we quantified spine measures in a subset of young adult Thy1-GFP-M mice (n= 5 dendritic segments from 3 males) taken from a separate motor cortex imaging experiment that was being run in the lab at the same time. These mice were also lightly restrained using the same restraint bags for 192s. The imaging relative to the timing of restraint was not identical to our main study (see supplementary figure 4) and is therefore presented as supplementary material. Analysis of that data using our Bayesian statistical model (see Supplementary table 3 for a summary of Bayes Factors), showed evidence in favour of the null hypothesis (i.e., post-restraint = baseline). This data does not replace a sham group but provides some data that suggests the restraint used in our study is unlikely to have caused the changes seen following rTMS.

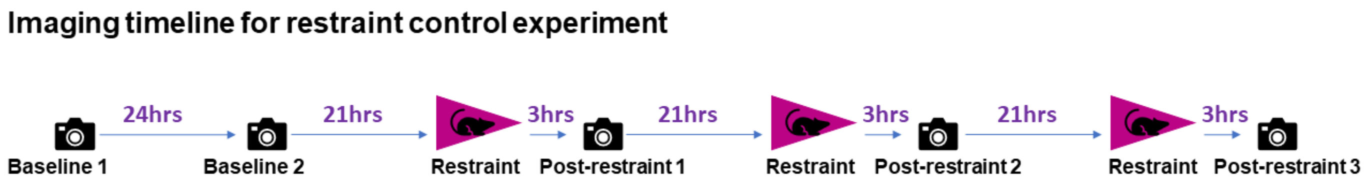

**Supplementary Figure 4. Imaging timeline of restraint only animals. NOTE: The timing of imaging relative to the restraint is not identical to the timeline in the main study (restraint + rTMS).**

**Supplementary Table 3. Summary of Bayes Factors for restraint data.**

| Spine Measure | Bayes Factors for null hypothesis (i.e., no difference between post-restraint and baseline) |
|---|---|
| **Density** | 15.4 |
| **Losses** | 5.2 |
| **Gains** | 4.0 |

### Supplementary material references

1. Dogdas B, Stout D, Chatziioannou AF, Leahy RM. Digimouse: a 3D whole body mouse atlas from CT and cryosection data. Phys Med Biol. 2007;52(3):577.

2. Yushkevich PA, Gao Y, Gerig G, editors. ITK-SNAP: An interactive tool for semi-automatic segmentation of multi-modality biomedical images. 2016 38th Annual

International Conference of the IEEE Engineering in Medicine and Biology Society (EMBC); 2016: IEEE.

9. Gasca F. Model-Based Improvements for Focal Transcranial Stimulation. Aus dem Institut fur Robotik und Kognitive Systeme. 2013.
